# How sample heterogeneity can obscure the signal of microbial interactions

**DOI:** 10.1101/520668

**Authors:** David W. Armitage, Stuart E. Jones

## Abstract

Microbial community data are commonly subjected to computational tools such as correlation networks, null models, and dynamic models, with the goal of identifying the ecological processes structuring microbial communities. Researchers applying these methods assume that the signs and magnitudes of species interactions and vital rates can be reliably parsed from observational data on species’ (relative) abundances. However, we contend that this assumption is violated when sample units contain any underlying spatial structure. Here, we show how three phenomena — Simpson’s paradox, context-dependence, and nonlinear averaging — can lead to erroneous conclusions about population parameters and species interactions when samples contain heterogeneous mixtures of populations or communities. At the root of this issue is the fundamental mismatch between the spatial scales of species interactions (micrometres) and those of typical microbial community samples (millimetres to centimetres). These issues can be overcome by measuring and accounting for spatial heterogeneity at very small scales, which will lead to more reliable inference of the ecological mechanisms structuring natural microbial communities.

## 1 Common “pattern-to-process” inferential methods yield erroneous results

Advances in sequencing technology offer microbiologists unprecedented access to the composition and dynamics of microbial communities [1]. Marker gene and metagenomic surveys regularly chronicle hundreds to thousands of taxa, many previously unknown, all seemingly co-occurring within their respective habitats. In possession of these large observational datasets, microbial ecologists have adapted theory and methods developed from plant and animal ecology to investigate how species interactions — such as competition, predation, and facilitation — structure microbial communities [2, 3].

Without experimental systems in which competition (or any other interaction) may be directly manipulated and detected, researchers often employ randomization-based null models, correlation networks, and population dynamic models to identify and quantify putative interspecific interactions from observational sequence data [4, 5, 6, 7]. Here, negative covariation between the abundances or relative abundance of taxa are commonly assumed to result from negative interspecific interactions such as competition. However, the utility of these methods for reliably parsing and quantifying signals of competition from alternative community assembly processes such as habitat filtering and trophic interactions has been disputed for decades [8].

Recently, a number of studies have challenged null model and correlation-based methods to recapitulate known interactions in well-studied marine intertidal habitats [9, 10, 11]. In all cases, these tests revealed troubling inaccuracies and discrepancies among the various methods, calling into question their ability to reliably identify true ecological interactions. For microbial communities, the only successful validations of these methods have occurred in simple, well-mixed liquid cultures [7]. Taken in concert, these studies highlight potential pitfalls in our ability to correctly identify species interactions when communities are sampled over underlying spatial heterogeneity. Most natural microbial communities are spatially structured and exhibit marked heterogeneity at multiple spatial scales. Failure to account for this underlying spatial heterogeneity in environmental samples can undermine our conclusions about the ecological processes structuring microbial assemblages [12].

## 2 Causes and consequences of heterogeneity in microbial samples

Typical sample volumes used for environmental marker gene and metagenomics studies are rarely smaller than 0.1 mL, but can be as large as 100 L of seawater and 100 g of soil in low-DNA habitats. Unless these samples come from a well-mixed, completely homogeneous medium, they will contain at least some amount of spatial structure. For example, a typical 0.25 g sample of soil containing particles 1 mm in diameter (i.e., a very coarse sand) will inevitably contain hundreds to thousands of discrete granules on which microbial communities can assemble. These discrete habitats can represent a heterogeneous array of environments or resources, each selecting for their own unique local microbial communities [13]. However, even a physicochemically homogeneous collection of particles can contain a mosaic of distinct microbial communities owing to the effects of limited or asymmetric dispersal, priority effects, and successional turnover.

Fine-scale heterogeneity in microbial communities appears to be a general property of environmental samples, having been repeatedly documented in aquatic, soil, fecal, leaf surface, and wastewater habitats [13, 14, 15, 16, 17, 18]. Owing to this, marker gene samples commonly represent a sum of sequence reads made over underlying environmental heterogeneity, leaving us with a bulk inventory of OTUs and their (often relative) abundances without their spatial context. Because microbial interactions such as resource competition, phage predation, DNA transfer, and syntrophy are hypothesized to take place at spatial scales much smaller than that of the typical bulk sample, it can be argued that many marker gene samples actually measure the *metacommunity* — a collection of semi-autonomous communities linked through dispersal [19]. In the following sections, we illustrate how collecting samples at the metacommunity scale can introduce errors into computational estimates of interspecific interactions by virtue of three phenomena: *Simpson’s paradox*, *context-dependence*, and *nonlinear averaging*. Note that although we present total abundance data throughout our scenarios, these phenomena also apply to compositional (i.e., relative abundance) data, which are more commonly collected in environmental marker gene surveys.

### 2.1 Simpson’s paradox

Simpson’s paradox refers to the reversal or negation of a statistical association between two variables, *X* and *Y*, when conditioned on a third variable, *Z* [20]. In ecology, this *Z* variable might include information on spatial variation among local patches, which, if accounted for, changes the direction of a trend at larger spatial scales [21]. Computational approaches to inferring microbial interactions can be sensitive to the effects of Simpson’s paradox. For instance, the inferred signs of interspecific correlation coefficients might change when comparing analytic results obtained from bulk community samples with results that have statistically accounted for underlying variation in microhabitats or resource availability within bulk samples.

To illustrate this point, consider a study that uses data obtained from bulk soil samples to infer the sign of interspecific interaction between two fungal taxa. If the true nature of this interaction is competitive, then our results are anticipated to reveal a negative correlation between the abundances of the two fungi. To add some realism to this scenario, let us assume that each of our samples represent collections of discrete microhabitats on which our focal taxa grow. Finally, we might also make the realistic assumption that both of our fungal taxa respond similarly to these discrete microhabitats such that sub-optimal habitats support fewer individuals of both species. If we populate bulk soil samples with random draws of simulated communities on each of three discrete microhabitat types (Fig. 1a), we find that even slight variation in the frequency distribution of these microhabitats within bulk samples leads to positive correlations between our two taxa, contradictory to their true, competitive local interactions. Furthermore, by repeating this experiment many times, each time re-assembling our bulk samples by populating them with equal numbers of randomly-selected discrete microhabitat particles, we encounter an overwhelming majority of cases where the inferred sign of interaction between our two taxa (positive) is the opposite of its true sign (negative) (Fig. 1b), leading us to erroneously conclude that these species are not strong competitors when, in truth, they are. Because of Simpson’s paradox, we contend that unless the assumption of homogeneity within and among microbial community samples is justified, interspecific interaction coefficients derived from correlation or model-based approaches should be interpreted with extreme caution, and should always include a statement concerning the spatial context of the sample including potential sources of underlying spatial heterogeneity.

**Figure 1:**
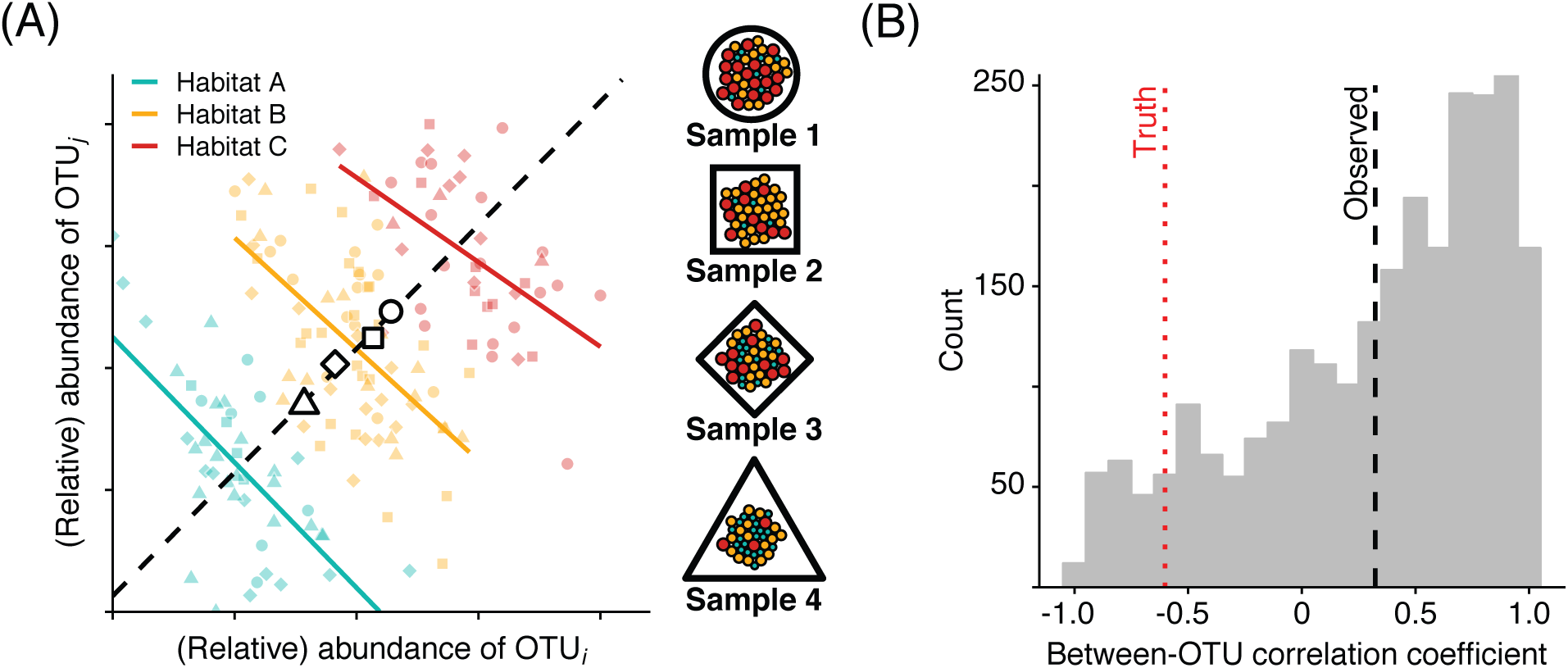
(A) Example of how Simpson’s paradox can influence the identification of interspecific interactions. Colored points show the abundances of OTUs *i* and *j* in samples across three discrete microhabitats. Though the OTUs compete with one another in all three habitat types, their population responses to each habitat are correlated. When bulk samples containing any variation in microhabitat composition are sequenced (denoted by white points), the inferred sign of species interactions can be erroneous. (B) A simulation analysis of 2500 individual OTU correlations taken from samples consisting of 250 randomly-assembled individual particles reveals that the average inferred sign of interspecific interactions is positive, whereas the true sign of these interactions (simulated at the scale of individual particles) is negative.

### 2.2 Context dependence

A common assumption of computational approaches for identifying species interactions is that the sign and strength of interactions are immutable across time and space. This assumption reduces the sample sizes required for estimating correlation coefficients or population parameters, and permits the use of graph theoretic descriptors of network structure (connectance, nestedness, etc.). However, numerous laboratory experiments have documented context-dependent interactions arising from variation in population densities, community composition, or environmental context, such that interactions measured at one place and time cannot reliably be extrapolated across habitats [22, 23, 24, 25]. For instance, a recent study documented predictable shifts in the sign of species interactions with changing resource concentrations in experimental yeast communities as cross-feeding gave way to competition [26] (Fig. 2a). The presence of predators can also mediate the sign of interspecific interactions through a variety of mechanisms [27] (e.g., Fig. 2b). Likewise, a meta-analysis of hundreds of experiments uncovered a strong effect of spatial heterogeneity on context-dependent species interactions [28]. Consequently, it is not unreasonable to expect the signs of microbial interactions to change across gradients of resource density, predation pressure, or other indicators of habitat quality (Fig. 2c). While temporal correlation network approaches might be used to circumvent the static interactions assumption at larger spatial scales or in well-mixed samples, they cannot account for variable interactions arising from underlying spatial heterogeneity within individual samples.

**Figure 2:**
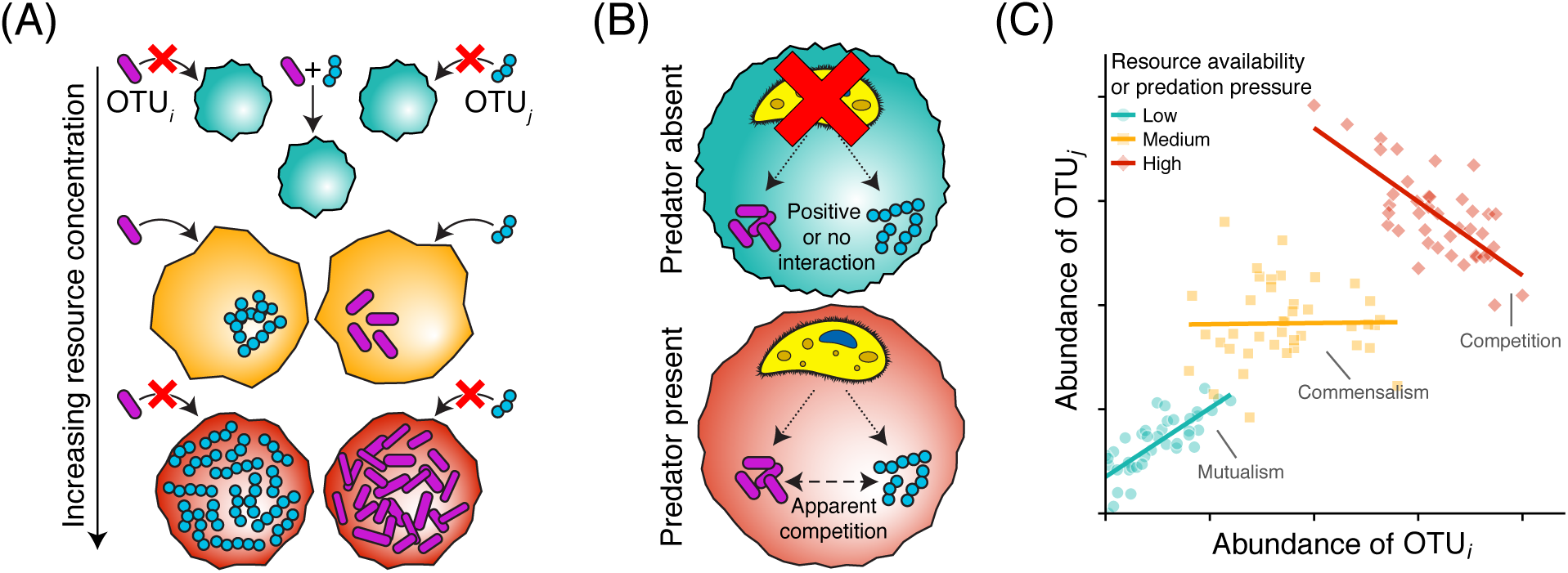
Examples of context-dependent species interactions. (A) Resource availability can modulate the sign of interspecific interactions. For instance, local resource limitation can weaken the strength of competition when (i) it selects for cross-feeding or another mutualistic, resource-concentrating behaviour, or (ii) when it limits the strength of interspecific negative density dependence. (B) Likewise, in situations where a shared predator is present, species that do not compete for shared resources can experience apparent competition by supplementing the predator densities. (C) These context-dependent interactions can lead to highly variable estimates of the signs of OTU interactions, depending on the spatial distribution of resources or predators within the sample.

From a theoretical perspective, context-dependence is hypothesized to be be a critical factor for maintaining diversity in spatially-structured communities [29]. For instance, the abilities of two competing microbial strains to coexist will be enhanced if the negative impacts of competition experienced by each strain are stronger in more favourable habitat patches [29]. Given that microbial species richness appears to peak in particulate, heterogeneous habitats (soil, sediments) [1], context-dependent interactions within these habitats may be quite common and important in promoting high levels of diversity. Currently, the extent of context-dependent interactions in spatially-structured microbial communities remains largely unknown. We note, though, that correlation network approaches have been successfully used to identify context-dependent interactions robust to experimental ground-truthing [30]. However, until the prevalence and magnitude of context-dependent microbial interactions are better understood, we encourage researchers to exercise caution when making general statements concerning any local estimates of interspecific interactions, ideally contextualizing results to the specific environment and scale at which measurements were taken.

### 2.3 Nonlinear averaging

The previous two sections concerned issues that arise when quantifying local microbial interactions from heterogeneous samples. However, we also face difficulties when using microbial community data collected at very small scales to quantify the aggregate behavior of aggregate microbial communities. Imagine that we are now able to obtain measurements of microbial populations at the scale of the individual microhabitat patches. Such data could be obtained, for instance, using a fluorescence in situ hybridization (FISH) approach to directly count cell densities on soil particles. Importantly, these data are collected at the spatial scale over which intraspecific interactions play out, which, in a heterogeneous sample experiencing dispersal among particles, is at the scale of individual microhabitat patches or particles. Called the *characteristic scale*, it is the scale which maximizes the ratio of deterministic signal to the influences of stochasticity and spatial heterogeneity [31], making it the optimal scale for measuring and characterizing the effects of deterministic species interactions.

Let us now envision a scenario where we wish to quantify whether a microbial OTU’s competitive ability is is a function of the local soil type. Since accurately estimating the strength of competition in our samples is of paramount importance, suppose we have conducted our sequencing surveys at appropriately small characteristic scales and have generated time series data from this assortment of individual particles. We then fit a population dynamic model to these data in order to estimate our OTU’s growth rate and competitive interactions among different soil types, adequately replicated within each type. The generalized Lotka-Volterra (gLV) population dynamic model is increasingly being utilized for this purpose. Fitting such a differential equation model requires estimating parameters describing a focal species’ growth rates and interspecific interactions. The gLV model commonly takes the form

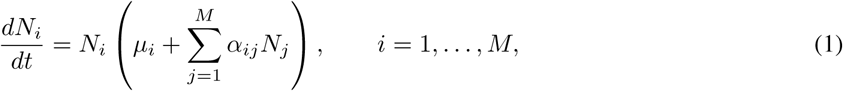

where *N*_*i*_ is the abundance of OTU *i*, *µ*_*i*_ is its maximum *per capita* growth rate, and *α*_*ij*_ is a parameter describing the proportional change in its growth rate with conspecific or heterospecific densities. Values of *α*_*ij*_ greater than zero imply that OTU *j* has a positive effect on OTU *i*, which might stem from interactions such as syntrophy, whereas values less than zero can signify interactions such as competition or chemical inhibition.

For illustrative purposes, let us simplify our problem of estimating competition among soil types by assuming that only our single focal OTU occupies our habitats, and so is only capable of experiencing intraspecific competition. This permits us to simplify our model to the case where (*i* = *j*), and define 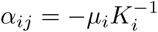, where *K*_*i*_ represents the local carrying capacity of our OTU *i*. This results in the familiar logistic population growth model describing decelerating microbial population growth with increasing population density. Expanding this model across a spatially-structured array of individual particles, we obtain the equation

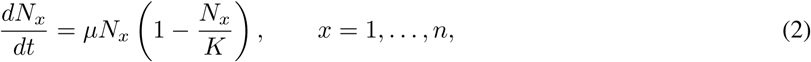

where *N*_*x*_ are the local sub-populations of our focal OTU on habitat particle *x*.

With a collection of population equations for our individual particles, we can now aggregate our local dynamics to obtain general growth parameters for our soil types. This scaling-up process requires a spatial averaging of local population dynamics. Crucially, because the average of a nonlinear function is not equal to the function of its averaged covariates (i.e., 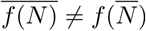), to scale up microbial population dynamics — which are almost unanimously nonlinear — by averaging across spatially-variable local populations will result in biases proportional to the spatial population variation and model’s nonlinearity. This principle, called *Jensen’s inequality*, has important consequences for our ability to accurately estimate scaled-up model parameters and make predictions from any gLV model fit to datasets containing underlying spatial heterogeneity.

The consequences of this spatial averaging process are illustrated in Fig. 3. For notational simplicity, we replace the growth function in equation 2, *µN*_*x*_(1 *− N*_*x*_/*K*), with *G*(*N*_*x*_). The spatially-averaged dynamical equation that we wish to obtain is 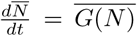. Calculating our population dynamic model using the spatial averages of the populations we have measured, 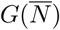, overestimates the correctly scaled-up population growth function, 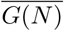. In Fig. 3c, we generated four collections of particles in which spatially-explicit populations have been randomly drawn from lognormal distributions having equal means but different variances (*σ*^2^). We then used these simulated data to fit four spatially-averaged population growth functions, 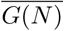. These results demonstrate how increasing the spatial variation among local populations has the effect of changing our scaled-up estimates of carrying capacity. The challenge for microbiologists is to accurately estimate 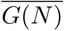 using our measured population densities, *N*_*x*_. Fortunately, if we have already collected these values, and if they can be reasonably fit to a population dynamic model, we can use the tools of *scale transition theory* [32, 33] to correctly obtain scaled-up population parameters. We briefly introduce these methods in the following section.

**Figure 3:**
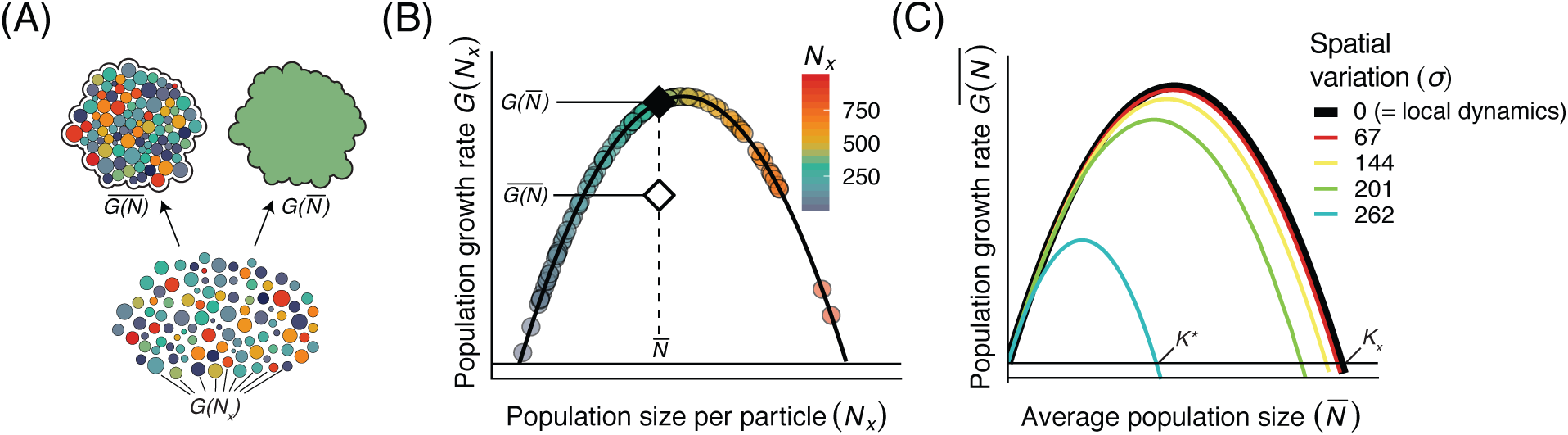
(A) Illustration of the concept of scaling-up local microbial community dynamics to quantify the behavior of an aggregate sample. Colors denote an OTU’s population sizes across a heterogeneous collection of particles governed by the shared, nonlinear dynamics, *G*(*N*_*x*_), shown in equation 2. Note the conceptual differences between aggregating these data by averaging over the local nonlinear dynamics, 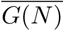, and by fitting our small-scale dynamical model to the average population density, 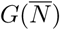. (B) The differences in these aggregation procedures result in differing estimates for scaled-up population dynamics. The black curve shows the logistic governing dynamics, *G*(*N*), of populations on individual particles (colored circles). Note the difference in growth rates between the correctly spatially-averaged growth function (white diamond) and growth function fit to the spatial average population density (black diamond). (C) Increasing the spatial variation of local populations results in vastly different spatially-averaged population dynamics. Here again, the black line denotes the local dynamics, *G*(*N*), which equals the the spatial average when there is no variation among subpopulations. For this concave-down function, increasing the spatial variation causes the scaled-up carrying capacity, *K*^∗^ to be smaller than the local carrying capacity, *K*_*x*_.

## 3 Recommendations moving forward

Despite the various ways in which spatial heterogeneity can subvert our interpretation or complicate our assessment of microbial community interactions and dynamics, we are optimistic that these issues can be surmounted with prudent data collection, analysis, and interpretation. The lurking effects of habitat heterogeneity are most effectively mitigated by quantifying microbial populations or communities at the spatial scales over which cell-cell interactions occur, which is on the scale of micrometers to millimeters. Sampling at this scale has successfully been accomplished using individual grains of sand [13], aquatic organic particles [34], and sludge granules [35] — all of which encountered marked heterogeneity among particles. Sampling at this scale is facilitated by technologies such as fluorescence-activated cell sorting and laser-assisted microdissection, which offer the opportunity to precisely and efficiently capture individual microscopic particles for sequencing. However, as we have seen, even measurements made at the appropriate characteristic scales can be challenging to generalize.

The restrictive assumptions of most correlation network and null models hinder our reliable assessment of microbial interactions in all but the most homogeneous samples. However, the influence of Simpson’s paradox and context-dependence may be surmounted by measuring and statistically accounting for the confounding effects of environmental and/or community variation among samples. Empirically, this might include increased efforts to quantify a sample’s micro-scale composition using spatially-resolved mass spectrometry and FISH techniques. Though challenging to collect, such data could then be used to more test the alternative hypotheses of habitat filtering and competition — both of which can feasibly manifest as identical community patterns in the presence of microhabitat variation.

While creative new statistical approaches for identifying nonlinear and context-dependent species interactions are becoming available [36], we suggest these methods be ground-truthed with more complex and realistic data than are currently in use. For example, rather than using time series simulated from equilibrial Lotka-Volterra equations to ground truth a new method, a more powerful validation routine could use data simulated from spatially-explicit agent-based models, which can test methods’ robustness to spatial heterogeneity, scale-dependence, and demographic stochasticity. We also encourage the inclusion of dynamic parameters in generalised Lotka-Volterra models. While it is challenging to estimate these parameters from observational data, experiments consistently show that microbial growth rate, carrying capacity, and interaction parameters are functions of their underlying environments. A benefit of including environmentally-dependent growth parameters in gLV models is that these models can then be used to quantify the effects of various coexistence-promoting mechanisms [29]. Context-dependent parameters also allow us to investigate the effects of environmental change on microbial populations and communities.

The increasing use of gLV models in microbial ecology also prompts us to account for the effects of nonlinear spatial averaging on scaled up population dynamics (section 2.3). Chesson’s scale transition theory [32, 33] provides a mathematical framework for tackling the issues of spatial heterogeneity and nonlinearities in gLV models. We introduce the scale transition using two simple models, but refer interested readers to the original papers for general scale transition approaches [32, 33]. Continuing from section 2.3, we can calculate the scaled-up population dynamics, 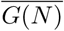, by accounting for the nonlinearity in *G*(*N*_*x*_) using its second derivative, *G*^′′^(*N*_*x*_), as well as the spatial variation in *N*_*x*_, measured by the spatial variance, Var(*N*). The full, spatially-averaged population model can be approximated as

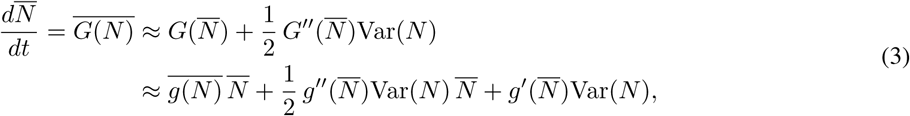

where 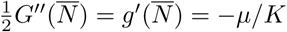. This approximation is exact when the growth function is quadratic (as is the case for logistic growth).

A similar, albeit more complicated scale transition can be calculated for a multispecies gLV model (eq. 1) [32]. This model is commonly used to identify interactions, denoted by the *α_ij_* parameters. By defining 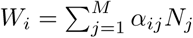 and *g*(*W*_*i*_) = *µ*_*i*_ +*W*_*i*_, the scaled up version of equation 1 can be written as a function of mean field terms, a nonlinearity term, and spatial variances and covariances:

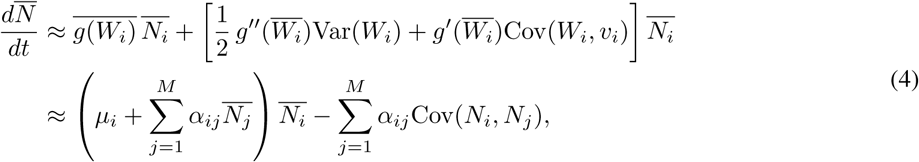

where 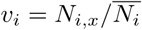. Once again, we see that the spatially-averaged population dynamics are not simply a function of average populations across space. However, the only extra information needed to calculate the scale transition are the spatial variances and covariances of the populations, which we can approximate by measuring local population densities across a sufficient number of particles within a sample. Thus, the calculation of scale transition terms is straightforward once they are defined for a particular dynamic model.

Given the potential for biases and errors stemming from the joint effects of underlying spatiotemporal heterogeneity and other methodological choices (e.g., relative abundance transformations, normalization techniques) [37], it may seem like the inference of species interactions from observational microbial data represents an *underdetermination* problem. That is, there may be multiple, or even infinite potential mechanisms capable of generating an observed community pattern. However, this problem, like many in ecology and evolution, can more precisely be described as an example of *contrast failure* [38]. Instead of a solution-free, underdetermined system, we instead have one where our failure to parse competing hypotheses is a transient consequence of data insufficiency. Access to better, more *contrastive* data, derived either experimentally or observationally at the appropriate spatiotemporal scales, will refine our ability to discriminate among alternative hypotheses. In the meantime, we do not advocate for the abandonment of ‘pattern-to-process’ approaches for deciphering microbial interactions. On the contrary, we are optimistic about continued methodological development in this area. In the meantime, we implore researchers to consider and confront the lurking effects of spatial structure on their inferred microbial interaction networks and growth parameters. At minimum, this could simply comprise a comment on the spatiotemporal scale over which the results are anticipated to hold and a description of the spatial structure contained within a sample unit.

